# Trophic position derived from amino-acid nitrogen isotopes reflect physiological status of both predator and prey over four decades

**DOI:** 10.1101/2021.01.25.428045

**Authors:** Agnes ML Karlson, Caroline Ek, Douglas Jones

## Abstract

Nitrogen isotope analyses of amino-acids (δ^15^N-AA) are increasingly used to decipher food webs. Interpretation of δ^15^N-AA in consumers relies on the assumption that physiological status has a negligible influence on the trophic enrichment factor (TEF). Recent experiments have shown that this is not always the case and there is a need to validate derived trophic position (TP) estimates using ecological data. We analyzed δ^15^N-AA in cod and herring (1980-2019) from the Baltic Sea, a species-poor system where dramatic reduction in condition status of cod has occurred. We expected that TEF_cod-herring_ in trophic AAs would increase during periods of poor cod growth, resulting in inflated TP estimates. We found that TEF and TP estimates were negatively linked to individual condition status, prey fat content and the hypoxic state of the ecosystem. Statistically adjusting for these variables resulted in lower cod TP, highlighting the importance of including ecological knowledge when interpreting TP.

**Scientific Significance Statement:** Nitrogen stable isotope analyses in amino acids are increasingly used in ecology to understand how environmental change impacts food-webs. Specifically, it is used to more accurately calculate trophic position (TP) of consumers. Controlled experiments have shown that physiological status can alter amino acid isotope composition and TP interpretation, but field studies are lacking. We use 40 years of archived material to demonstrate that TP estimates in Baltic Sea cod and its prey herring are directly related to physiological status. This has important implications for interpreting the real trophic ecology of consumers under environmental stress. By simultaneously measuring condition status in both predator and prey it is possible to adjust for them as confounding variables and decipher actual consumer TP.

## Introduction

Evaluating the impacts of environmental change on marine ecosystems and implementing sustainable management practices requires an improved understanding of the trophodynamics of food webs. Trophic positioning (TP) of consumers is instrumental when assessing food web structure (Post 2002) and biomagnification of contaminants (Rolff et al. 1993). Stable isotope ratios of nitrogen (^15^N:^14^N), expressed as δ^15^N (parts per thousand differences from a standard), are commonly used to estimate TP in consumers, since the heavy isotope is enriched with each trophic level in a rather predictable way i.e. the difference in permille between consumer and diet (e.g. Minagawa and Wada 1984; McCutchan et al. 2003). However, this so-called trophic enrichment factor (TEF) is known to increase in starving or slow-growing animals (e.g. Martinez del Rio et al. 2005). Furthermore, the δ^15^N baseline, which depends on the isotopically distinct nitrogen sources used by primary producers, varies considerably both in space and time. Hence, interpreting δ^15^N in bulk samples, particularly for higher trophic levels is highly challenging (Post 2002).

Analysis of δ^15^N in specific amino acids (AA) in consumers has become increasingly popular because it allows for TP and baseline δ^15^N estimates, without sampling the base of the food web (Chikaraishi et al. 2009, Ohkouchi et al. 2017). Trophic AAs increase in δ^15^N in consumers due to transamination, whereas source AAs show no or very little change in their δ^15^N-values (McClelland and Montoya 2002, Chikaraishi et al. 2009). δ^15^N in AAs can potentially provide a more accurate TP estimation in food web studies than δ^15^N in bulk material (Blanke et al. 2017, Bowes et al. 2015). It is however a developing technique and the problem with variable TEFs remains. Laboratory experiments testing how δ^15^N-AAs in fish respond to diets of varying protein quantity and quality often find conflicting results regarding the derived TEF estimates, where some AAs seem to be more sensitive to diet quality than others (e.g. McMahon et al. 2015, Nuche-Pascual et al. 2018). How estimates of TP based on AAs are affected in real world ecosystems where individuals can adapt their behavior (move, diet shift, compensatory feeding) is unknown.

Here we take advantage of the uniquely long time series of environmental monitoring data and archived samples from the Baltic Sea (Reisch et al 2018) to link physiological status in both cod and its prey herring with measured isotope data. During the last four decades this ecosystem has gone from a cod-dominated system to a forage fish dominated system, largely as a consequence of overfishing and increased hypoxia (e.g. Möllman and Diekman 2012). Increased competition for zooplankton among sprat and herring contributed to decreased condition of herring (e.g. Casini et al 2010), but with improvements in the last two decades (Karlson et al. 2020). For cod, a decrease by around 30% in mean body condition and sharp declines in growth (Casini et al. 2016, Mion et al. 2020) coincide with the rapid increase in hypoxic water volume since the mid 90s (Savchuk 2018). Stomach analyses suggest no change in diet for larger cod in recent decades (Neuenfeldt et al. 2020), although increased compensatory feeding due to poor diet quality has been proposed (Svedäng et al. 2020). Hypoxia is likely to influence young cod by limiting the amount of benthos (Neuenfeldt et al. 2020), but for larger cod the direct physiological effects from hypoxia exposure on metabolism and condition status (Limburg and Casini 2018), is likely more important.

The well documented trends in cod and herring condition status over the last 4 decades has likely influenced their δ^15^N values. Low growth rates and poor food conversion efficiency could lead to catabolism in trophic AAs (and hence higher isotope discrimination) to provide energy for basal metabolism and in turn, inflate TP estimates (e.g. McMahon et al. 2015). Environmental stress such as hypoxia could also cause elevated δ^15^N values from altered metabolic pathways (Popuin et al. 2014) leading to enrichment in trophic AAs such as glutamic acid (Ek et al. 2018). We expect that over time, trophic AAs and derived TEF and TP estimates, in both herring and cod, are confounded by the physiological status of the fish (Fig 1). More specifically, low condition factor (i.e. Fulton’s K, Ricker 1975) and fat content is expected to result in higher TEF and TP from growth rate dependent isotope discrimination. Additionally, cod δ^15^N values and TP estimates are expected to be influenced by the condition/fat of its prey (Fig 1) and finally, hypoxia-induced isotope discrimination effects are also expected for cod. We test the influence of physiological status and environmental stress on AA-δ^15^N in cod and herring muscle tissue collected over the last 40 years using various regression analyses.

**Fig 1.**
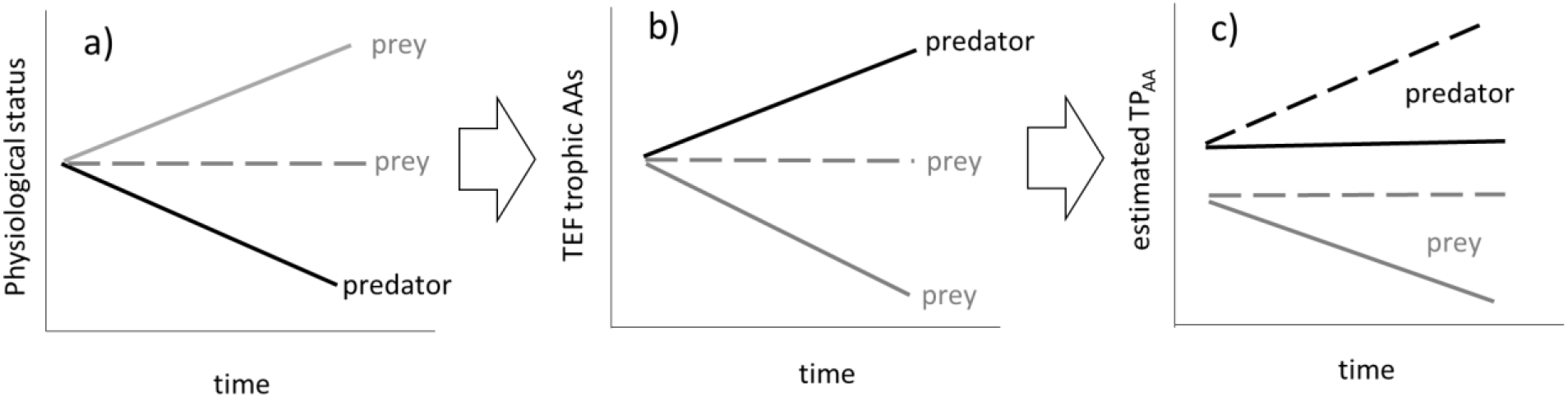
Temporal development in prey and predator a) physiological status, declining for predator while unchanged or improving for prey, b) resultant expected trophic enrichment factor (TEF) for the trophic amino-acids and c) resultant estimated TP for prey and predator, where predatorTP is dependent not only on its own TEF but also on the TEF in prey during the two scenarios; solid and dotted lines for predators should be compared to the development of prey with solid and dotted lines, respectively.

## Methods

### Sample preparation

Samples of cod and herring were provided by the Environmental Specimen Bank (ESB, long term frozen storage, −30 °C) at the Swedish Museum of Natural History. Cod were caught at a monitoring station, south east of the island of Gotland (56° 53’N, 18° 38’E). Fish were 35.3 cm ± 3.6 (total length) with an estimated age (otolith reading) of 3 years; age determination in Baltic cod is highly uncertain, especially in recent years (Hüssy et al. 2019). Herring (total length 18.3 ± 1.1 cm, age 3-5) were caught outside the island Landsort (58° 42’N, 18° 04’E), in the western Gotland basin. All fish were caught during the autumn (October-November) and frozen immediately after capture. Cod in this size class feeds almost entirely on pelagic prey (Neuenfeldt et al. 2020) and herring is mainly zooplanktivorous (Möllman and Köster 2004). We use herring as a proxy for pelagic forage fish prey since this is the species available, but importantly herring constitute the main proportion of cod diet based on weight (for cod in this size class the contribution of herring in diet is unchanged over time) (Neuenfeldt et al. 2020).

Ten individuals from each species and year, with as equal sex ratio as possible, over a 40-year period (cod; 1981-2019, one missing year 2014, herring 1980-2018; missing years 1982, 1993, 1998, 2000, 2006, 2013 and 2015) were sampled for isotope analyses. White muscle tissue from above the lateral line were dissected (epidermis and fat tissue were removed; Pinnegar and Polunin 1999). The samples were freeze-dried and homogenized whereafter a pool of the material based on the ten individuals for each year was used for AA-δ^15^N analyses; 3 mg of each pooled sample was weighed in glass vials. In addition, bulk analyses of δ^13^C (and δ^15^N) were analysed (1 mg sample in tin capsules). δ^13^C values show only a slight change relative to the diet and is therefore used to link consumers to the primary producers at the base of the food chain (e.g. δ^13^C in fish can be compared to the δ^13^C baseline of blue mussels, Hanson et al. 2020). All samples were sent to the UC Davis Stable Isotope Facility, US, for analysis. δ^13^C values were adjusted for the C:N ratio according to Post et al. (2007). Data and metadata will be made available in the dryad data repository.

### Data analyses and Statistics

Biological data on length, weight, age and fat content (muscle fat for herring and liver fat for cod, expressed as % of wet weight; Karlson et al. 2020) were compiled from the ESB database (publicly available at www.sgu.se) and annual mean values were calculated for use as potential explanatory variables in statistical analyses (see below). Fulton’s condition index (based on length and weight; Ricker 1975) was calculated from the individuals used for isotope measurements. Fat data was in 1981-1996 based on 20 individuals, 1997-2014 on 10 individuals and since 2015 a mean value of two pools of 12 individuals (comparable over time according to Bignert et al. 2014).

Trophic position based on AA-δ^15^N was calculated for both cod and herring according to the equations by Chikaraishi et al. (2009) using equation 1 and Nielsen et al. (2015) using eq. 2.

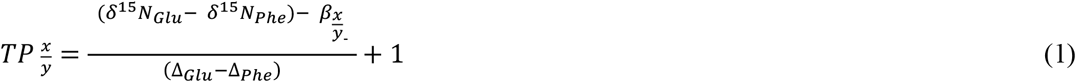

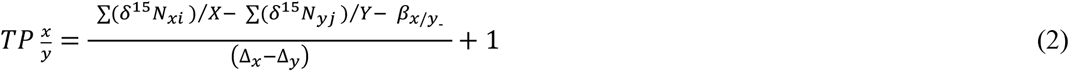

where N_xi_ are the δ^15^N values of trophic AA_*i*_, and N_yj_ are the δ^15^N values of source AAj. The letters in subscript _*i*_ and _*j*_ corresponds to the different trophic and source AAs respectively, in the equation. β_x/y_ corresponds to the difference between the δ^15^N values of trophic AAs (x) and source AAs (y) in primary producers, Δ_x_ and Δ_y_ are the ^15^N trophic enrichment factors (TEF) for each AA(s) _x_ and _y_, respectively. Values for β_x/y_ and TEF differ between the equations; in eq. 1 TP was calculated from Glu-Phe using the following values for β_x/y_ and TEF (3.4 and 7.6). In eq. 2, the values were 2.9 for β_x/y_ and 5.9 for TEF (the AA Met was omitted similar to Nielsen et al. 2015).

Pearson correlations were performed to test which of the source AA δ^15^N values covaried over time for cod and herring (i.e. δ^15^N-Phe_cod_ with δ^15^N-Phe_herr_); this was done to validate the assumption of a stable predator-prey relationship between the trophic groups. We explored if a 1 year lag on the cod data would improve correlations. Yearly estimates of trophic enrichment factor for cod (TEF) in both source and trophic AAs was calculated as the difference between cod and herring δ^15^N in each AA (cod-herring δ^15^N values) for each year.

We carried out statistical breakpoint analyses in R (R Core Team, 2019) using the strucchange package (Kleiber et al 2002) to compute breakpoints in regression relationships of fat% and Fulton’s K over time in both species. Additionally, we performed breakpoint analyses for the isotope data (bulk δ^13^C and bulk δ^15^N, δ^15^N in the individual AAs, TEF_cod-herring_ values and TP values). The optimal number, and corresponding year(s), of breakpoints in a time series were selected based on BIC estimates. Unidirectional time trends in physiological status, isotope data and derived trophic metrics were assessed by non-parametric Mann-Kendall tests in time series based on identified breakpoints. During the deterioration period, potential negative relationship between fat% (arcsine transformed) and Fulton’s K and each AAs TEF_cod-herring_ were tested using Pearson correlations.

Generalized linear models (GLMs) were used to test the effects of physiology, potential diet change over time (bulk δ^13^C), prey condition (fat% in herring, best proxy for diet quality, Karlson et al. 2020) and environmental stress (hypoxic volume for the Baltic proper; integrated yearly values until 2016 extracted from Savchuk 2018) on TP estimates (eq 1 and 2). For simplicity, fat% was used as physiological condition for cod and the interaction between cod and herring fat% was included (in initial models without the interaction term, fat% was a better predictor than Fulton’s K). For cod each time period (defined using breakpoint analyses) was analysed separately since different physiological effects were expected during these periods (e.g. Casini et al. 2016); these time periods also coincide with the known Baltic Sea regime shift (e.g. Möllman et al. 2009). Herring had no clear physiological breakpoint (see results), and the entire time series was analysed, with Fulton, fat% and the δ^13^C baselines used as initial predictors. Final models were the best combination of predictors based on parsimony and AIC scores. Analyses were performed in Statistica in the GLZ module using a log-link (residuals were visually inspected for normality).

To reduce the amount of unexplained variation and hence increase the statistical power to detect true time trends in TP (Fig 1), potential confounding factors for the TP estimates (the variables selected in the final GLM models based on AIC criteria) were adjusted for by multiple regression (Kleinbaum et al. 2008, Ek et al. 2021) using the software package PIA (Bignert 2007). This analysis estimates the TP values as if the covariates were constant at their average values. Breakpoint analyses and time trends using Kendall-tau test were performed on adjusted TP values and compared to non-adjusted results.

## Results

### Temporal changes in physiological status, isotope data and trophic metrics

Fulton’s K and fat% correlated positively and significantly for both species (cod; r=0.47, p<0.003, herring r=0.63, p<0.001, Fig. 2). According to breakpoint analyses fat% and Fulton’s K in cod began decreasing in 1993 and 1995 respectively. Both condition metrics decreased after 1993 (Fulton; tau=−0.63, p<0.001, fat% tau=−0.53, p<0.001) and we defined this year as the start of a deteriorating period for cod in subsequent analyses. For herring, breakpoints were identified at 1990 (fat%, increase after 1990, tau=0.30, p=0.050) and 2010 (Fulton’s K, decrease until 2010, tau=−0.34, p=0.017), the entire time series were used when exploring herring physiological status in relation to its TP values.

**Figure 2.**
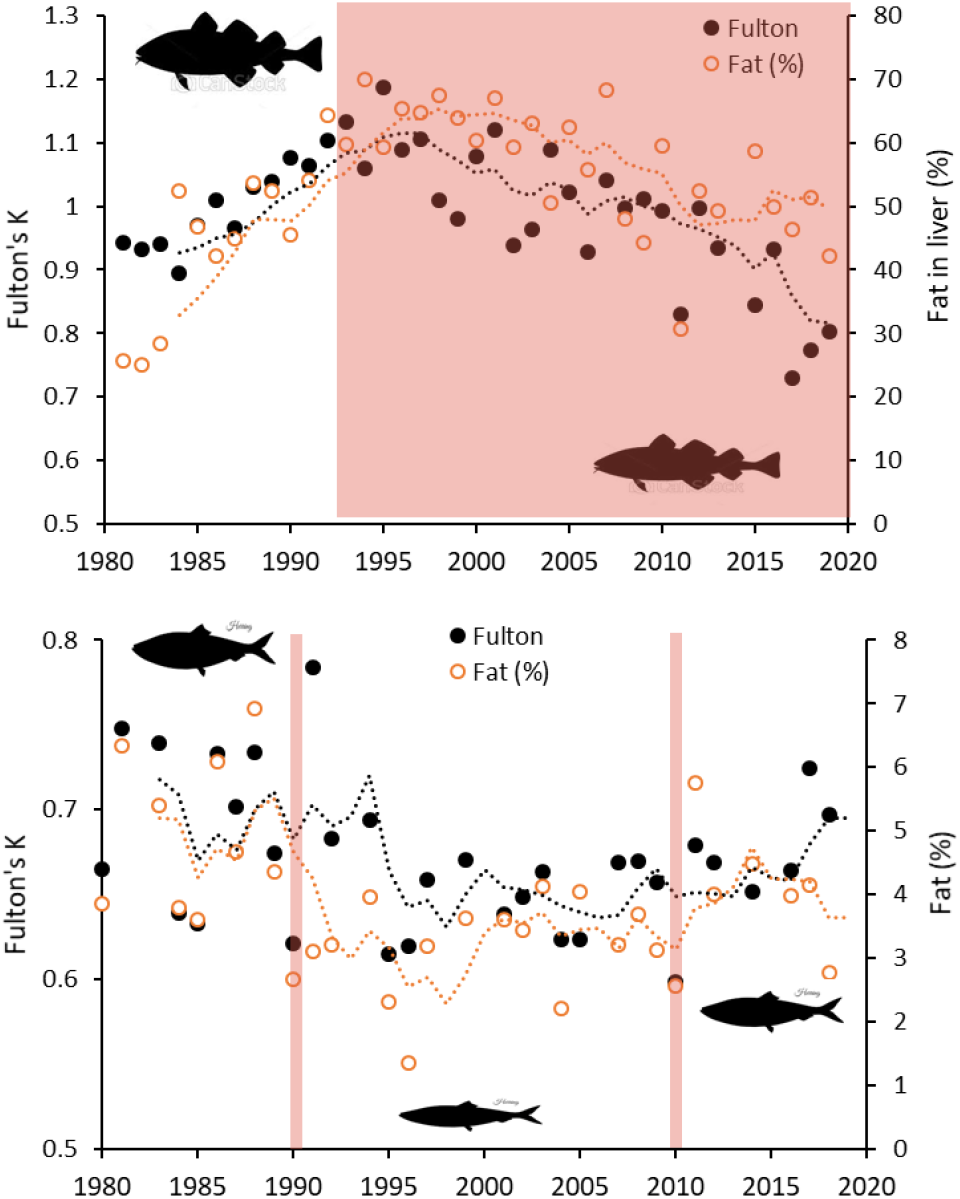
Physiological status in cod (top) and herring (bottom) 1980-2019 (dotted lines are five year moving averages). Red area defines the start of the cod deterioration period and the red lines for herring denote shifts in fat% (1990, followed by improvement, Table 1) and Fulton’s condition index (2010, Table 2). Shifts are based on breakpoint analyses (see text for details).

Bulk δ^15^N (Fig 3a) correlated significantly positively to all trophic AAs in both species (on average r=0.8), and to most of the source AAs but with lower r-values (on average 0.2 for cod and 0.5 for herring, Table S1-S2). For the entire time series, δ^15^N in the source AAs phenylalanine (Phe) and methionine (Met) as well as the trophic AA aspartine (Asp) covaried significantly positively in herring and cod (Pearson’s r=0.4-0.5, Fig 3d, Table S3). Correlations with a 1 year lag for cod improved correlation coefficients for these AAs during the deterioration period (e.g. Phecod-herring from r=0.28, p<0.25 to r=0.68, p<0.001). δ^13^C in both cod and herring covaried with the δ^13^C blue mussel baseline over the entire time series (cod; r=0.23, p<0.06, herring; r=0.30, p<0.09, cod; Fig 3b) and demonstrated three similar breakpoints, occurring once per decade (1992-1993, 2003-2006, and 2010-2013).

**Figure 3.**
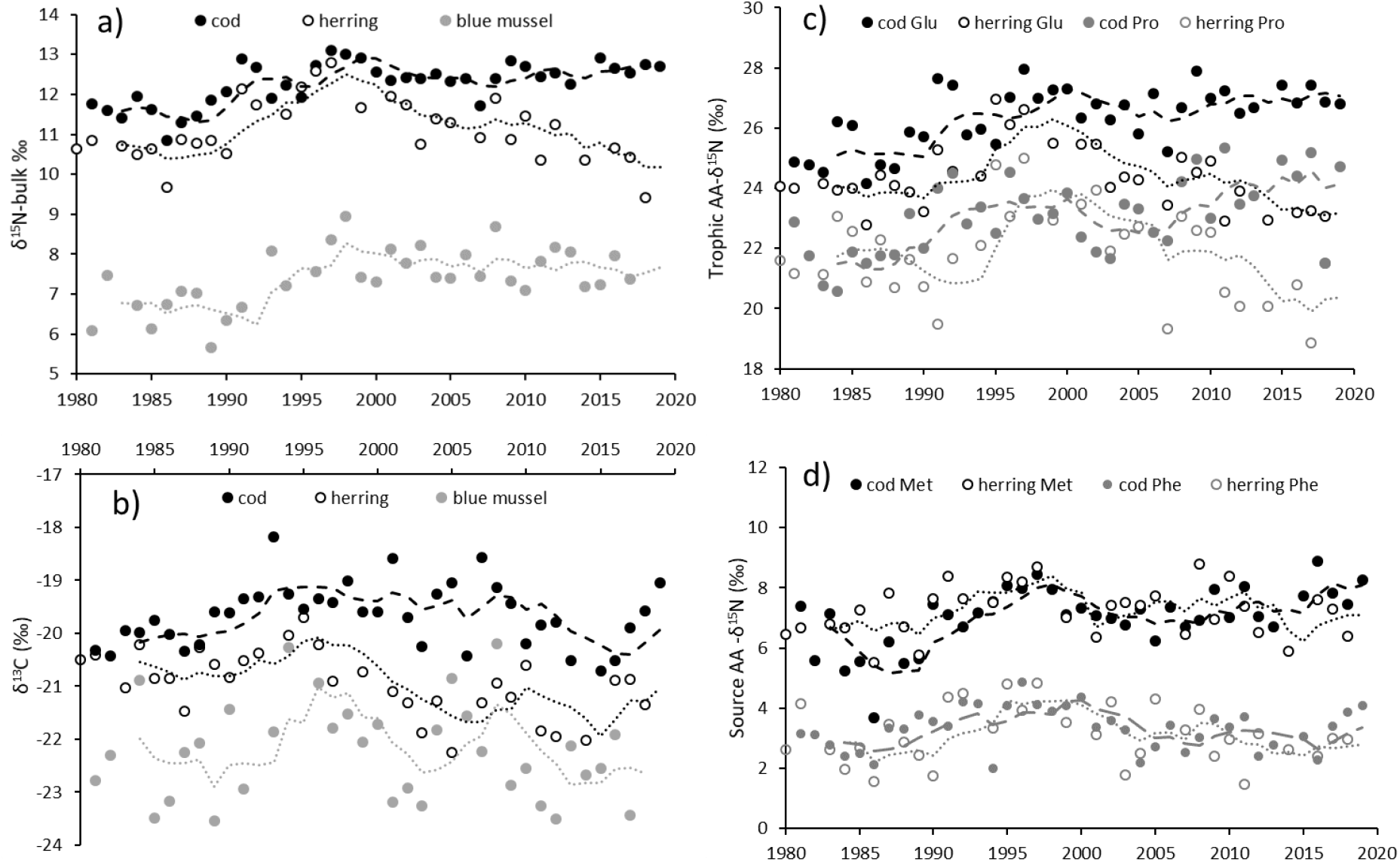
Long-term patterns in bulk-δ15N and bulk-δ13C for cod, herring and blue mussels (left panel, a and b) and in AA-δ15N in cod and herring for the trophic AAs c) Glutamic acid and Proline and the source AAs d) Phenylalanine and Methionine. Glu and Pro correlate highly with all other trophic AAs, for both species (Table S2). Dotted lines are five year moving averages.

Breakpoints for δ^15^N-AAs in herring were found in 1990 (Ala, Val, Phe, Lys, Gly, Ser, His and Tyr, bulk-δ^15^N) or in 1994 (Glu, Asp, Pro, Ile, Leu). Cod had a highly variable pattern regarding δ^15^N breakpoints (Table S4), however the calculated TEF_cod-herring_ breakpoints were found in 1994 for several trophic AAs (Table S4), whereafter significant unidirectional increases followed (Table S5). The breakpoint(s) for the TP estimates in cod and herring (Fig 4) and the potential trends as assessed from Mann-Kendall tests differed between the two methods (Table 1).

**Figure 4.**
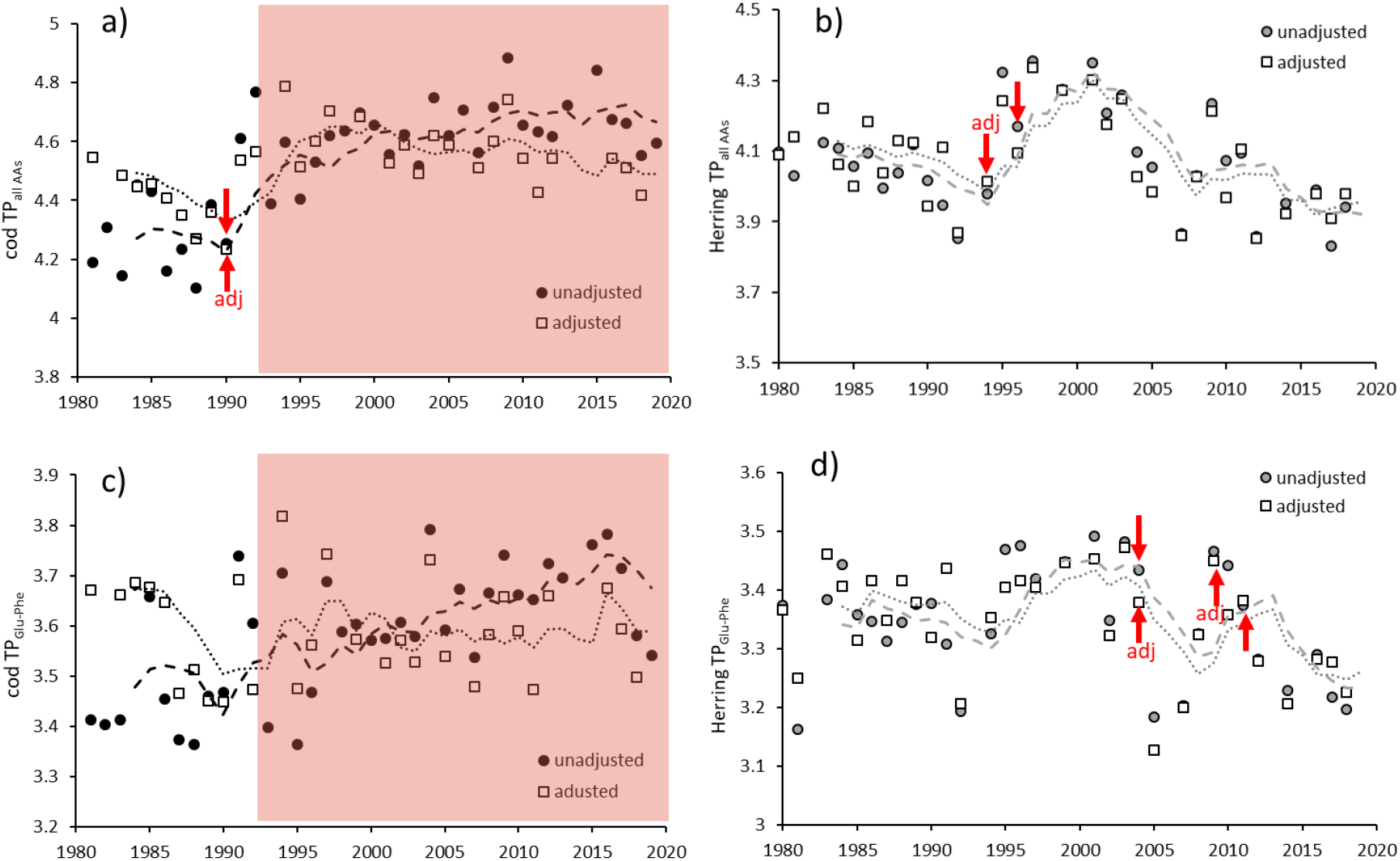
Long-term patterns in trophic position (TP) of cod (left panel) and herring (right panel) calculated using all AAs (a and b) and only Glu and Phe (c and d). TP is shown with and without statistical adjustments (see text); red arrows demonstrate breakpoints followed by increases for unadjusted TP in cod (Table 1) or decreases for unadjusted herring TP_all_ (increase until 2004 for both unadjusted and adjusted TP_Glu-Phe_). Note that the adjusted TP_all_ for cod decreases significantly after the breakpoint (1990) and during the deterioration period (from 1993, red shaded area). Statistical results can be found in Table 1. Dotted lines are five year moving averages.

**Table 1.** TP derived using the two methods (denoted by TP_glu-phe_ and TP_all_) for herring (1980-2018) and cod (time series divided into before and after the deterioration period starting in 1993, see text) was used in Generalized linear models (GLM) to test the effects of physiological condition, diet and hypoxia (see Statistics section). Best models based on AIC and the estimates of these variables are shown and then used as covariates in subsequent multiple regression to calculate adjusted TP values. CV (coefficient of variation), breakpoint year and trends (Man-Kendall-tests) are shown for both non-adjusted and adjusted TP values. Bold values denote a significant trend (p<0.035 according to Bonferroni correction of pairwise comparisons before/after adjustment; if Bonferroni correction is applied across all 15 comparisons performed, p<0.014). Trends for entire or combined (cod) time series are tested before and after the breakpoint(s) with an additional adjustment for hypoxia for cod (see also Fig 4).

| Response variable | GLM models based on AIC |  |  |  |  | Log-linear multiple regression |  | Breakpoint (BP) |  | Mann-Kendall trend before / after TP BP |  | Mann-Kendall trend after cod fat BP (1993) |  |
| --- | --- | --- | --- | --- | --- | --- | --- | --- | --- | --- | --- | --- | --- |
|  | Best model | Est. | SE | Wald | p | CV un-adj. | CV <u>Adjusted</u> | un-adj. | <u>Adjusted</u> | tau (p-value) un-adj. | tau (p-value) <u>Adjusted</u> | tau (p-value) un-adj. | tau (p-value) <u>Adjusted</u> |
| Herring TP<sub>glu-phe</sub> entire period | fulton's K | -0.31 | 0.11 | 7.89 | 0.005 | 2.9 | 2.6 | 2004, 2011 | 2004, 2009 | 0.31 (0.05) / -0.73 (0.06) | 0.21 (0.20) / **-0.64 (0.035)** | **-0.53 (<0.01)** | **-0.46 (<0.01)** |
| Herring TP<sub>all</sub> entire period | fulton's K | -0.29 | 0.14 | 4.27 | 0.039 | 3.5 | 3.1 | 1996 | 1994 | -0.14 (0.49) / **-0.67 (<0.01)** | -0.39 (0.07) / **-0.46 (<0.01)** | **-0.49 (0.010)** | **-0.46 (<0.01)** |
| Cod TP<sub>glu-phe</sub> after 1993 | cod fat | -0.54 | 0.16 | 11.39 | <0.001 | 2.7 | 2.6 | NA | NA | NA | NA | 0.27 (0.07) | 0.14 (0.44) |
|  | herring fat | -5.57 | 2.17 | 6.58 | 0.010 |  |  |  |  |  |  |  |  |
|  | cod fat*herr fat | 10.79 | 3.71 | 8.45 | 0.004 |  |  |  |  |  |  |  |  |
| Cod TP<sub>glu-phe</sub> before 1993 | herring fat | -1.84 | 0.58 | 10.23 | 0.001 | 3.8 | 2.3 | NA | NA | NA | NA | *before 1993:* 0.13 (0.58) | *before 1993:* **-0.67 (<0.01)** |
| Cod TP<sub>all</sub> after 1993 | cod fat | -0.41 | 0.12 | 12.46 | <0.001 | 2.1 | 2.0 | NA | NA | NA | NA | 0.21 (0.14) | -0.08 (0.68) |
|  | herring fat | -4.50 | 1.58 | 8.08 | 0.005 |  |  |  |  |  |  |  |  |
|  | cod fat*herr fat | 7.45 | 2.70 | 7.63 | 0.006 |  |  |  |  |  |  |  |  |
| Cod TP<sub>all</sub> before 1993 | cod fat | 0.94 | 0.13 | 55.67 | <0.001 | 4.2 | 2.7 | NA | NA | NA | NA | *before 1993:* 0.31 (0.16) | *before 1993:* -0.31 (0.22) |
|  | herring fat | 5.96 | 1.27 | 21.88 | <0.001 |  |  |  |  |  |  |  |  |
|  | cod fat*herr fat | -15.06 | 2.42 | 38.62 | <0.001 |  |  |  |  |  |  |  |  |
| Cod TP entire period |  |  |  |  |  | Fat-adj. | <u>Hypox. adj</u> | un-adj. | Fat-adj. / <u>Hypox-adj</u> | un-adj. | <u>Hypox-adj</u> | Fat-adj, see above | <u>Hypox-adj</u> |
| Cod TP<sub>glu-phe</sub> | NA | NA | NA | NA | NA | 3.1 | 2.8 | no BP | 1990 / no BP | **0.39 (<0.01)** | -0.01 (0.99) | 0.27 (see above) | 0.01 (1) |
| Cod TP<sub>all</sub> | NA | NA | NA | NA | NA | 3.0 | 2.7 | 1990 | 1990 / 1990 | -0.06 (0.92) / 0.26 (0.05) | **-0.79 (<0.01)** / -0.29 (0.07) | 0.21 (see above) | **-0.39 (0.02)** |

### Linking consumer physiology to isotope data and trophic metrics: adjustments of time trends

Fat% and Fulton’s K in cod was significantly correlated to TEFs for trophic AAs but not to source AAs during its deterioration period (Table S6). Herring TP (both methods) was best explained by its condition (Fulton), while cod TP (both methods) was best explained by both herring fat and its own fat content as well as their interaction (Table 1). Although the time series before the deterioration period is quite short, the model resulted in the same selection of variables, but with opposite effects (+ vs −) compared to the deterioration period (Table 1). Adjusting TP estimates for condition variables resulted in reduced variation in all time series (CV, Table 1) and after Bonferroni corrections a change in trend was found for half of the comparisons (i.e. unadjusted vs adjusted), with examples of both trends being detected and trends disappearing after adjustment (Table 1, Fig 4, Fig S1). In particular changes were detected for TP_glu-phe_ estimates, e.g. the increasing trend in the entire time series for unadjusted TP_glu-phe_ in cod disappeared after adjustment. The adjusted TP data in cod correlated significantly with hypoxic volume (Fig S2, TP_all_ r=0.72, TP_glu-phe_ r=0.56, p<0.01 in both cases, corresponding values for unadjusted TP estimates were r=0.44 p<0.02 and r=0.33, p<0.10). Hypoxic volume was initially not selected in the best model for unadjusted TP, but after first fat and then hypoxia adjustment, the time trend during the deterioration period changed to negative for TP_all_ (Fig 4a, Table 1). For herring, the breakpoints were detected 2 years earlier after adjustment and a decreasing trend was detected after the last breakpoint in TP_glu-phe_ (Table 1, Fig 4b and d).

## Discussion

This study demonstrates long-term divergent changes in trophic position (TP) for cod and its prey herring, as assessed from the amino-acid nitrogen isotope composition (δ^15^N-AA). As hypothesized (Fig 1), the calculated trophic enrichment factor (TEF) and the TP values derived from the Chikaraishi et al. (2009) and Nielsen et al. (2015) equations were influenced by physiological status. Adjusting for physiology detrended the previously increasing trend for TP in cod using both methods.

As expected, negative correlations were established between TEFs for trophic AAs (but not source AAs) and physiological/condition status in cod during the period of poor cod growth (1993-2018). Accordingly, the decreasing fat content explained increasing TP in cod during this period. For herring, the morphometric (Fulton) condition index best explained TP over the entire time period (the lower condition the higher the TP estimate). It is expected that slow growth (from limited or bad quality food) results in elevated δ^15^N-values due to larger isotope discrimination (e.g. excretory losses will be comparatively high for slow growing animals not incorporating nitrogen to body mass). However, cod TP was also influenced by herrings improving fat content. The improved condition status of its prey in recent decades, with its lower δ^15^N-values (due to reduced isotope discrimination during faster growth), partly counteracted the inflated δ^15^N-values in cod resulting in a less pronounced increase in its TP (Fig 1). Considering the dramatic decrease in cod condition, higher TP estimates would likely have been observed if herrings condition status (and δ^15^N) were stable over time. There are indications of food limitations for Baltic cod (Casini et al. 2016). Under low food availability, the additional effect of metabolic turnover rates on endogenous N leads to higher fractionation and TP estimates; an effect that was only visible in strongly increasing TEF estimates in trophic AAs from the early 1990s when cod show a clear change in condition. Our study shows that prey physiology effects on isotope composition are propagated up the food web. Further support for the influence of herring condition (rather than a change in diet) on isotope composition in cod come from the significant correlation between herring fat% and cod δ^13^C (Fig S3); lipids are depleted in δ^13^C, and at times when herring are in good condition status this results in lower cod δ^13^C values (despite the C:N correction).

In addition to growth-related effects, exposure to hypoxic conditions likely also influence isotope discrimination and TP estimates. During hypoxic conditions both direct and indirect effects can be expected. Exposure to toxic hydrogen sulphide can result in physiological costs from combating toxic exposure (Calow 1991) and result in increased ^15^N in trophic AAs (Ek et al 2018). In addition, indirect effects on cod metabolism can occur due to migration to warmer, more oxygen-rich waters (Claireaux and Chabot 2016) possibly leading to altered ^15^N fractionation pattern (Poupin et al 2014). It is hypothesized that (small) cod are exposed to hypoxic conditions whilst foraging after benthos (Limburg & Casini 2019, Brander 2020). We note that Baltic cod spawning size has decreased over time (Köster et al 2017) and exposure to hypoxia in the fish in this study (~35 cm) could be a result of summer spawning migrations (up to several months in deep, saline but oxygen poor water; Nielsen et al. 2013). The stronger correlation between TP-estimates and hypoxia after fat-adjustment of TP (Fig S2) strongly suggests that both growth status and environmental stress influence TEF and TP estimates in this species (Table 1). We have not focused on interpreting the absolute TEF or TP estimates but note that different methods for TP calculation result in differences of at least 0.5 units. The fat and hypoxia adjustments had a more pronounced effect on TP calculated from the AAs Glu and Phe compared to the method using all AAs (Nielson et al. 2015). After adjustment using the Nielsen method a breakpoint in 1990 was still evident which may reflect patterns in fat and several AAs for herring. Further studies are required to investigate which method of TP calculation provide the most reliable TP estimates

To conclude, great care should be taken when reconstructing food webs from times series of archived samples based on δ^15^N values in AAs if there is no information about consumer condition status and prey δ^15^N and condition status. This study demonstrates the importance of including physiological status metrics, not only in predators but also in their prey, to avoid erroneous interpretations of TP in food web studies.

## Author contributions

AK initiated the study, acquired funding and wrote the first draft of the paper. CE and DJ contributed equally to the study by commenting on the study approach and writing. All authors conducted statistical analyses.

## Acknowledgements

The retrospective isotope analyses in cod was funded by Naturvårdsverket (Swedish EPA) and specifically Elisabeth Nyberg to AK (contract 213-19-016), and in herring with funding from the Stockholm University Baltic Sea Centre to AK. All fish have over the decades been sampled within the Swedish contaminant monitoring program, commissioned by SEPA and stored at the Environmental Specimen Bank at the Swedish Museum of Natural History (SMNH). We thank the laboratory staff at SMNH especially Per-Arvid Berglund for dissecting the fish. Victoria Engström and Sara Forsberg prepared all the fish samples for isotope analyses and Chris Yarnes at UC Davis performed the amino acid isotope analyses. We thank Matias Ledesma for discussions on TP calculations and Sture Hansson, Carl Rolff and Stefan Neuenfeldt for discussions on herring and cod feeding ecology.

## Supplementary material

Table S1-S2. Correlations of bulk δ^15^N and δ^15^N-AAs for cod (Table S1) and herring (Table S2).

Table S3. Correlations between δ^15^N-AAs in cod and herring.

Table S4. Breakpoints for δ^15^N in AAs and TEFs for cod.

Table S5. Mann-Kendall trends for individual TEFs in AAs.

Table S6. Correlations among TEFs and physiological status, before and after the cod deterioration period

Fig S1. Trophic position in cod with fat adjustment only.

Fig S2. Correlations between trophic position in cod, with and without fat-adjustment, and hypoxic volume.

Fig S3. Correlation between herring fat% and cod δ^13^C in the entire time series.

## References

Bignert, A., 2007. PIA statistical application developed for use by the Arctic Monitoring and Assessment Programme (available from www.amap.no).

Bignert, A., Eriksson, U., Nyberg, E., Miller, A. and S. Danielsson. 2014. Consequences of using pooled versus individual samples for designing environmental monitoring sampling strategies. Chemosphere 94: doi: 10.1016/j.chemosphere.2013.09.096

Blanke, C. M., Chikaraishi, Y., Takizawa, Y., Steffan, S. A., Dharampal, P. S. and J. M. Vander Zanden. 2017. Comparing compound-specific and bulk stable nitrogen isotope trophic discrimination factors across multiple freshwater fish species and diets. Canadian J Fish Aquat Sci 74: 1291–1297, doi.org/10.1139/cjfas-2016-0420

Brander, K. 2020. Reduced growth in Baltic Sea cod may be due to mild hypoxia. ICES J Mar Sci 77: 2003–2005, doi.org/10.1093/icesjms/fsaa041.

Bowes, R. and J. Thorp. 2015. Consequences of employing amino acid vs. bulk‐tissue, stable isotope analysis: a laboratory trophic position experiment. Ecosphere 6: 1–12.

Calow, P. 1991. Physiological costs of combating chemical toxicants: ecological implications. Comp. Biochem. Phys. C. 100: 3–6.

Casini, M., V. Bartolino, J. C. Moliniero, and G. Kornilovs. 2010. Linking fisheries, trophic interactions and climate: Threshold dynamics drive herring *Clupea harengus* growth in the central Baltic Sea. Mar Ecol Prog Ser 413: 241–252.

Chikaraishi, Y., Ogawa, N. O., Kashiyama, Y., Takano, Y., Suga, H., Tomitani, A., Miyashita, H., Kitazato, H. and N. Ohkouchi. 2009. Determination of aquatic food-web structure based on compound specific nitrogen isotopic composition of amino acids. Limnol. Oceanogr: Methods 7: 740–750.

Claireaux, G. and D. Chabot. 2016. Responses by fishes to environmental hypoxia: integration through Fry’s concept of aerobic metabolic scope. J Fish Biol. 88: 232–251.

Ek, C., Holmstrand, H., Mustajärvi, L., Garbaras, A., Bariseviciue R. T., Sapolaite, J., Sobek, A., Gorokhova, E. and A. M. L. Karlson. 2018. Using compound-specific and bulk stable isotope analysis for trophic positioning of bivalves in contaminated Baltic Sea Sediments. Environ. Sci Technol. 52: 4861–4868.

Hanson, N., Larsson, Å., Parkkonen, J., Faxneld, S., Nyberg, E., Bignert, A., Ek, H. H., Bryhn, A., Olsson, J., Karlson, A. M. L. and L. Förlin. 2020. Ecological changes as a plausible explanation for differences in uptake of contaminants between European perch and eelpout in a coastal area of the Baltic Sea. Environ. Toxicol. Pharmacol. 80: 103455 doi.org/10.1016/j.etap.2020.103455

Hüssy, K., Radtke K., Plikshs, M., Oeberst, R., Baranova, T., Krumme, U., Sjöberg, R., Walther, Y. and H. Mosegaard. 2016. Challenging ICES age estimation protocols: Lessons learned from the eastern Baltic cod stock. ICES J Mar Sci 73: 2138–2149 doi.org/10.1093/icesjms/fsw107

Karlson AML, Gorokhova E, Gårdmark A, Pekcan-Hekim Z, Casini M, Albertsson J, Sundelin B, Karlsson O, Bergström L. 2020. Linking consumer physiological status to food-web structure and prey food value in the Baltic Sea. AMBIO 49: 391–406doi:10.1007/s13280-019-01201-1

Kleiber, C., Hornik, K., Leisch, F., and Zeileis, A. 2002. strucchange: An r package for testing for structural change in linear regression models. J Stat Softw. 7: 1–38.

Kleinbaum, D.G., Kupper, L.L., Nizam, A., Muller, K.E., 2008. Applied Regression Analysis and Other Multivariate methods Fourth edition. Brooks Cole.

Köster, F. W., Huwer, B., Hinrichsen, H. H., Neumann, V., Makarchouk, A., Eero, M., Dewitz, B. V., Hüssy, K., Tomkiewicz, J., Margonski, P., Temming, A., Hermann, J. P., Oesterwind, D., Dierking, J., Kotterba, P. and M. Plikshs. 2017. Eastern Baltic cod recruitment revisited—dynamics and impacting factors. ICES J Mar Sci 74: 3–19

Limburg, K. E., and M. Casini. 2019. Otolith chemistry indicates recent worsened Baltic cod condition is linked to hypoxia exposure. Biology Letters 15: 20190352.

Limburg, K. E., and M. Casini. 2018. Effect of marine hypoxia on Baltic sea cod Gadus morhua: evidence from otolith chemical proxies. Front. Mar. Sci. 5: 482.

Martínez del Rio, C. and B. O. Wolf. 2005. Mass-balance models for animal isotopic ecology. In Physiological and Ecological Adaptations to Feeding in Vertebrates (Starck, M. J., Wang, T., Eds.). Science Publishers: Enfield, New Hampshire, chap. 6, 141–174 pp.

McCutchan, J. H., Lewis, W. M., Kendall, C. and C. C. McGrath. 2003. Variation in trophic shift for stable isotope ratios of carbon, nitrogen, and sulfur. Oikos 102: 378–390.

McClelland, J. W and J. P. Montoya. 2002. Trophic relationships and the nitrogen isotopic composition of amino acids in plankton. Ecology 83: 2173–2180.

McMahon, K. W., Thorrold, S. R., Elsdon, T. S., and M. D. McCarthy. 2015. Trophic discrimination of nitrogen stable isotopes in amino acids varies with diet quality in a marine fish. Limnol. Oceanogr. 60: 1076–1087. doi.org/10.1002/lno.10081

Mion, M., Haase, S., Hemmer‐Hansen, J., Hilvarsson, A., Hüssy, K., Krüger‐Johnsen, M., Krumme, U., McQueen, K., Plikshs, M., Radtke, K., Schade, F. M., Vitale, F. and M. Casini. 2020. Multidecadal changes in fish growth rates estimated from tagging data: A case study from the Eastern Baltic cod (*Gadus morhua*, Gadidae). Fish Fish. doi.org/10.1111/faf.12527

Minagawa, M. and E. Wada. 1984. Stepwise enrichment of 15N along food chains: Further evidence

Möllmann, C., Kornilovs, G., Fetter, M., and F. W. Köster. 2004. Feeding ecology of central Baltic Sea herring and sprat. J Fish Biol. 65: 1563–1581.

Möllmann, C., Diekmann, R., Müller‐Karulis, B., Kornilovs, G., Plikshs, M., and P. Axe. 2009. Reorganization of a large marine ecosystem due to atmospheric and anthropogenic pressure: a discontinuous regime shift in the Central Baltic Sea. Glob Change Biol. 15: 1377–1393, doi: 10.1111/j.1365-2486.2008.01814.x

Neuenfeldt, S., Bartolino, V., Orio, A., Andersen, K. H., Andersen, N. G., Niiranen, S., Bergström, U., Ustups, D., Kulatska, N., and Casini, M. 2020. Feeding and growth of Atlantic cod (*Gadus morhua* L.) in the eastern Baltic Sea under environmental change. ICES J Mar Sci. 77: 624–632. doi:10.1093/icesjms/fsz224

Nielsen, J. M., Popp, B. N. and M. Winder 2015. Meta-analysis of amino acid stable nitrogen isotope ratios for estimating trophic position in marine organisms. Oecologia 178: 631–642.

Nielsen, B., Hüssy K., Neuenfeldt, S., Tomkiewicz, J., Behrens, J. W. and K. H. Andersen 2013. Individual behaviour of Baltic cod *Gadus morhua* in relation to sex and reproductive state. Aquat. Biol. 18: 197–207.

Nuche-Pascual, M. T., Lazo, J. P., Ruiz-Cooley, R. I., and S. Z. Herzka. 2018. Amino acid-specific δ^15^N trophic enrichment factors in fish fed with formulated diets varying in protein quantity and quality. Ecol. Evol. doi: 10.1002/ece3.4295

Ohkouchi, N., Chikaraishi, Y., Close, H. G., Fry, B., Larsen, T., Madigan, D. J., McCarthy, M. D., McMahon, K. W., Nagata, T. and Y. I. Naito. 2017. Advances in the application of amino acid nitrogen isotopic analysis in ecological and biogeochemical studies. Org. Geochem. 113: 150–174.

Pinnegar, J. K. and N. V. C. Polunin, 1999. Differential fractionation of δ^13^C and δ^15^N among fish tissues: implication for the study of trophic interactions. Funct. Ecol 13: 225–231

Post, D. M. 2002. Using stable isotopes to estimate trophic position: Models, methods, and assumptions. Ecology 83: 703–718.

Post, D. M., Layman, C. A., Arrington, D. A., Takimoto, G., Quattrochi, J. and C. G. Montana. 2007. Getting to the fat of the matter: Models, methods and assumptions for dealing with lipids in stable isotope analyses. Oecologia 152: 179–189.

Poupin, N., Mariotti, F., Huneau, J. F., Hermier, D. and H. Fouillet. 2014. Natural isotopic signatures of variations in body nitrogen fluxes: a compartmental model analysis. PLoS Comput Biol. 10 e1003865.

R Core Team. 2019. R: A language and environment for statistical computing. R Foundation for Statistical Computing, Vienna, Austria. URL https://www.R-project.org/

Reusch, T. B. H., and others. 2018. The Baltic Sea as a time machine for the future coastal ocean. Sci. Adv. 4: eaar8195. doi:10.1126/sciadv.aar8195

Ricker, W. E. 1975. Computation and interpretation of biological statistics of fish populations. Bull. Fish. Res. Bd. Can. 191: 1–382.

Rolff, C., Broman, D., Näf, C., and Y. Zebühr. 1993. Potential biomagnification of PCDD/FS – new possibilities for quantitative assessment using stable isotope trophic position. Chemosphere 27: 461–468

Savchuk, O. P. 2018. Large-Scale Nutrient Dynamics in the Baltic Sea, 1970-2016. Front. Mar. Sci. doi.org/10.3389/fmars.2018.00095

Svedäng, H., Thunel,l V., Pålsson, A., Wikström S. A., and M. J. Whitehouse. 2020. Compensatory Feeding in Eastern Baltic Cod (*Gadus morhua*): Recent Shifts in Otolith Growth and Nitrogen Content Suggest Unprecedented Metabolic Changes. Front. Mar. Sci. doi.org/10.3389/fmars.2020.00565

