## Supplementary material for "Trophic position derived from amino-acid nitrogen isotopes reflect physiological status of both predator and prey over four decades"

**Table S1**. Pearson correlations among δ^15^N in trophic (Glu to Leu) and source (Phe to Thr) amino acids and bulk δ^15^N in **cod** over the entire time series. Values are Pearson coefficient (r) and red denote p<0.05.

|  | Glu | Asp | Ala | Val | Pro | Ile | Leu | Phe | Lys | Gly | Ser | Met | His | Tyr | Thr |
| --- | --- | --- | --- | --- | --- | --- | --- | --- | --- | --- | --- | --- | --- | --- | --- |
| Glu |  | 0.93 | 0.85 | 0.84 | 0.68 | 0.72 | 0.94 | 0.39 | 0.73 | -0.37 | 0.19 | 0.71 | 0.33 | 0.61 | -0.80 |
| Asp | 0.93 |  | 0.89 | 0.81 | 0.61 | 0.66 | 0.89 | 0.33 | 0.70 | -0.40 | 0.19 | 0.69 | 0.29 | 0.55 | -0.71 |
| Ala | 0.85 | 0.89 |  | 0.70 | 0.44 | 0.59 | 0.83 | 0.33 | 0.68 | -0.21 | 0.31 | 0.62 | 0.29 | 0.58 | -0.66 |
| Val | 0.84 | 0.81 | 0.70 |  | 0.76 | 0.58 | 0.90 | 0.38 | 0.66 | -0.30 | 0.23 | 0.83 | 0.44 | 0.79 | -0.72 |
| Pro | 0.68 | 0.61 | 0.44 | 0.76 |  | 0.44 | 0.72 | 0.25 | 0.40 | -0.25 | 0.16 | 0.70 | 0.30 | 0.56 | -0.62 |
| Ile | 0.72 | 0.66 | 0.59 | 0.58 | 0.44 |  | 0.61 | 0.20 | 0.56 | -0.62 | -0.33 | 0.51 | -0.07 | 0.61 | -0.78 |
| Leu | 0.94 | 0.89 | 0.83 | 0.90 | 0.72 | 0.61 |  | 0.40 | 0.70 | -0.26 | 0.28 | 0.73 | 0.35 | 0.64 | -0.75 |
| Phe | 0.39 | 0.33 | 0.33 | 0.38 | 0.25 | 0.20 | 0.40 |  | 0.46 | 0.14 | 0.42 | 0.37 | 0.36 | 0.41 | -0.07 |
| Lys | 0.73 | 0.70 | 0.68 | 0.66 | 0.40 | 0.56 | 0.70 | 0.46 |  | -0.18 | 0.24 | 0.42 | 0.16 | 0.53 | -0.47 |
| Gly | -0.37 | -0.40 | -0.21 | -0.30 | -0.25 | -0.62 | -0.26 | 0.14 | -0.18 |  | 0.65 | -0.24 | 0.20 | -0.29 | 0.62 |
| Ser | 0.19 | 0.19 | 0.31 | 0.23 | 0.16 | -0.33 | 0.28 | 0.42 | 0.24 | 0.65 |  | 0.18 | 0.49 | 0.03 | 0.14 |
| Met | 0.71 | 0.69 | 0.62 | 0.83 | 0.70 | 0.51 | 0.73 | 0.37 | 0.42 | -0.24 | 0.18 |  | 0.38 | 0.72 | -0.56 |
| His | 0.33 | 0.29 | 0.29 | 0.44 | 0.30 | -0.07 | 0.35 | 0.36 | 0.16 | 0.20 | 0.49 | 0.38 |  | 0.32 | -0.12 |
| Tyr | 0.61 | 0.55 | 0.58 | 0.79 | 0.56 | 0.61 | 0.64 | 0.41 | 0.53 | -0.29 | 0.03 | 0.72 | 0.32 |  | -0.60 |
| Thr | -0.80 | -0.71 | -0.66 | -0.72 | -0.62 | -0.78 | -0.75 | -0.07 | -0.47 | 0.62 | 0.14 | -0.56 | -0.12 | -0.60 |  |
| bulk δ^15^N | 0.92 | 0.93 | 0.83 | 0.82 | 0.65 | 0.74 | 0.87 | 0.38 | 0.68 | -0.38 | 0.17 | 0.79 | 0.28 | 0.64 | -0.71 |

**Table S2**. Pearson correlations among δ^15^N in trophic (Glu to Leu) and source (Phe to Thr) amino acids and bulk δ^15^N in **herring** over the entire time series. Values are Pearson coefficient (r) and red denote p<0.05.

|  | Glu | | Asp | Ala | Val | Pro | Ile | Leu | Phe | Lys | Gly | Ser | Met | His | Tyr | Thr |
| --- | --- | --- | --- | --- | --- | --- | --- | --- | --- | --- | --- | --- | --- | --- | --- | --- |
| Glu | |  | 0.97 | 0.80 | 0.68 | 0.85 | 0.77 | 0.95 | 0.70 | 0.46 | 0.31 | 0.44 | 0.53 | 0.61 | 0.64 | 0.16 |
| Asp | | 0.97 |  | 0.79 | 0.62 | 0.82 | 0.81 | 0.94 | 0.68 | 0.48 | 0.26 | 0.40 | 0.53 | 0.62 | 0.59 | 0.04 |
| Ala | | 0.80 | 0.79 |  | 0.46 | 0.61 | 0.60 | 0.71 | 0.65 | 0.43 | 0.46 | 0.52 | 0.42 | 0.60 | 0.50 | 0.00 |
| Val | | 0.68 | 0.62 | 0.46 |  | 0.64 | 0.31 | 0.75 | 0.49 | 0.18 | 0.32 | 0.50 | 0.48 | 0.44 | 0.30 | 0.25 |
| Pro | | 0.85 | 0.82 | 0.61 | 0.64 |  | 0.72 | 0.84 | 0.46 | 0.17 | 0.26 | 0.21 | 0.46 | 0.41 | 0.32 | 0.11 |
| Ile | | 0.77 | 0.81 | 0.60 | 0.31 | 0.72 |  | 0.73 | 0.40 | 0.25 | 0.13 | 0.02 | 0.13 | 0.45 | 0.23 | -0.11 |
| Leu | | 0.95 | 0.94 | 0.71 | 0.75 | 0.84 | 0.73 |  | 0.62 | 0.38 | 0.36 | 0.49 | 0.53 | 0.55 | 0.52 | 0.12 |
| Phe | | 0.70 | 0.68 | 0.65 | 0.49 | 0.46 | 0.40 | 0.62 |  | 0.68 | 0.26 | 0.57 | 0.43 | 0.73 | 0.60 | 0.28 |
| Lys | | 0.46 | 0.48 | 0.43 | 0.18 | 0.17 | 0.25 | 0.38 | 0.68 |  | 0.09 | 0.27 | 0.55 | 0.56 | 0.60 | 0.19 |
| Gly | | 0.31 | 0.26 | 0.46 | 0.32 | 0.26 | 0.13 | 0.36 | 0.26 | 0.09 |  | 0.67 | 0.19 | 0.25 | 0.13 | 0.35 |
| Ser | | 0.44 | 0.40 | 0.52 | 0.50 | 0.21 | 0.02 | 0.49 | 0.57 | 0.27 | 0.67 |  | 0.38 | 0.49 | 0.50 | 0.44 |
| Met | | 0.53 | 0.53 | 0.42 | 0.48 | 0.46 | 0.13 | 0.53 | 0.43 | 0.55 | 0.19 | 0.38 |  | 0.29 | 0.55 | 0.35 |
| His | | 0.61 | 0.62 | 0.60 | 0.44 | 0.41 | 0.45 | 0.55 | 0.73 | 0.56 | 0.25 | 0.49 | 0.29 |  | 0.57 | 0.09 |
| Tyr | | 0.64 | 0.59 | 0.50 | 0.30 | 0.32 | 0.23 | 0.52 | 0.60 | 0.60 | 0.13 | 0.50 | 0.55 | 0.57 |  | 0.32 |
| Thr | | 0.16 | 0.04 | 0.00 | 0.25 | 0.11 | -0.11 | 0.12 | 0.28 | 0.19 | 0.35 | 0.44 | 0.35 | 0.09 | 0.32 |  |
| bulk δ ^15^N | | 0.94 | 0.91 | 0.75 | 0.58 | 0.72 | 0.66 | 0.91 | 0.75 | 0.61 | 0.30 | 0.50 | 0.59 | 0.60 | 0.75 | 0.16 |

**Table S3**. Correlations between δ^15^N in AAs in cod and herring over the entire time series (1981-2018). Values are Pearson coefficient (r) and red denote p<0.05. Values in bold are the pairwise comparisons of main interest. Correaltions with cod lag 1 year during its deterioration period (1993 and onwards) improve correlation coefficients (data not shown).

|  |  | |  | |  | |  | | | **Herring AAs** | | | |  | | |  |  | | |  | |  | |
| --- | --- | --- | --- | --- | --- | --- | --- | --- | --- | --- | --- | --- | --- | --- | --- | --- | --- | --- | --- | --- | --- | --- | --- | --- |
| **Cod AAs** | Glu | Asp | | Ala | | Val | | Pro | Ile | | Leu | Phe | Lys | | Gly | Ser | | | Met | His | | Tyr | | Thr |
| Glu | **0.30** | 0.28 | | 0.26 | | 0.17 | | 0.16 | 0.11 | | 0.26 | 0.20 | 0.31 | | -0.09 | 0.18 | | | 0.44 | 0.02 | | 0.43 | | 0.04 |
| Asp | 0.41 | **0.42** | | 0.39 | | 0.25 | | 0.25 | 0.27 | | 0.38 | 0.24 | 0.34 | | -0.01 | 0.21 | | | 0.51 | 0.07 | | 0.47 | | 0.02 |
| Ala | 0.37 | 0.37 | | **0.20** | | 0.28 | | 0.31 | 0.18 | | 0.36 | 0.19 | 0.21 | | -0.14 | 0.11 | | | 0.37 | -0.01 | | 0.40 | | -0.01 |
| Val | 0.31 | 0.28 | | 0.46 | | **0.08** | | 0.15 | 0.13 | | 0.23 | 0.29 | 0.36 | | 0.01 | 0.22 | | | 0.50 | 0.06 | | 0.43 | | -0.03 |
| Pro | 0.11 | 0.08 | | 0.19 | | 0.04 | | **-0.14** | 0.03 | | 0.11 | 0.21 | 0.43 | | -0.07 | 0.21 | | | 0.39 | 0.01 | | 0.41 | | 0.12 |
| Ile | 0.21 | 0.20 | | 0.29 | | -0.05 | | 0.22 | **0.16** | | 0.15 | 0.12 | 0.23 | | -0.04 | 0.08 | | | 0.15 | -0.06 | | 0.25 | | -0.27 |
| Leu | 0.36 | 0.31 | | 0.35 | | 0.19 | | 0.26 | 0.15 | | **0.30** | 0.28 | 0.33 | | -0.06 | 0.22 | | | 0.52 | 0.06 | | 0.47 | | 0.10 |
| Phe | 0.51 | 0.48 | | 0.38 | | 0.26 | | 0.31 | 0.39 | | 0.39 | **0.38** | 0.31 | | -0.12 | 0.03 | | | 0.31 | 0.10 | | 0.31 | | 0.19 |
| Lys | 0.27 | 0.25 | | 0.21 | | 0.08 | | 0.23 | 0.23 | | 0.18 | 0.07 | **0.19** | | -0.36 | -0.18 | | | 0.22 | 0.09 | | 0.31 | | -0.19 |
| Gly | -0.14 | -0.13 | | -0.27 | | 0.04 | | -0.19 | -0.10 | | -0.08 | -0.14 | -0.31 | | **-0.18** | -0.22 | | | -0.21 | 0.04 | | -0.23 | | 0.19 |
| Ser | 0.19 | 0.16 | | -0.02 | | 0.36 | | -0.08 | 0.05 | | 0.24 | 0.23 | 0.24 | | -0.17 | **0.04** | | | 0.31 | 0.01 | | 0.21 | | 0.25 |
| Met | 0.31 | 0.33 | | 0.39 | | 0.06 | | 0.10 | 0.19 | | 0.27 | 0.29 | 0.34 | | 0.07 | 0.22 | | | **0.52** | -0.04 | | 0.38 | | -0.03 |
| His | 0.10 | 0.14 | | 0.06 | | 0.13 | | -0.09 | 0.12 | | 0.15 | -0.02 | -0.05 | | -0.06 | 0.08 | | | 0.28 | **-0.17** | | 0.12 | | 0.08 |
| Tyr | 0.26 | 0.27 | | 0.38 | | -0.04 | | 0.22 | 0.26 | | 0.16 | 0.15 | 0.19 | | -0.15 | -0.12 | | | 0.25 | -0.03 | | **0.20** | | -0.31 |
| Thr | -0.22 | -0.18 | | -0.29 | | -0.18 | | -0.14 | 0.00 | | -0.17 | -0.24 | -0.40 | | -0.10 | -0.28 | | | -0.38 | -0.02 | | -0.38 | | **0.06** |

**Table S4.** Statistical breakpoint(s) in time series of δ^15^N-AA in cod and for AA-TEF_cod-herring_. Bulk δ^15^N in cod had a breakpoint at 1988, (same year for blue mussel; 1990 for bulk δ^15^N herring) and for TEF_cod-herring_ in bulk δ^15^N there was a breakpoint in1994.

| AA | | δ^15^N in cod AAs | δ^15^N-AA TEF_cod-herring_ |
| --- | --- | --- | --- |
| Glu | | 1985, 1990, 1995, 2007 | 1994 |
| Asp | | 1985,1992, 1997, 2008, 2013 | 1994 |
| Ala | | 1985, 1992, 1997, 2000 | 1992, 2004 |
| Val | | no BP | 1988 |
| Pro | | no BP | 1994 |
| Ile | | 1988, 1996 | no BP |
| Leu | | 2000 | no BP |
| Phe | | no BP | no BP |
| Lys | | 2003 | 1990 |
| Gly | | 2005, 2013 | no BP |
| Ser | | no BP | no BP |
| Met | | 1986, 1999 | 1987 |
| His | | no BP | no BP |
| Tyr | | 1988 | no BP |
| Thr | | 1990 | 1990 |

**Table S5**. Unidirectional trends in AA-TEF_cod-herring_ after breakpoint (second breakpoint for Ala, 2004) or entire time series if there was no breakpoint (compare Table S4). Values in bold are lower than the Bonferroni corrected (n=15 AAs) p-value of 0.013; please note that time series length varies depending on breakpoint year (if any breakpoint).

| AA | tau value | p-value |
| --- | --- | --- |
| Glu | **0.67** | **<0.001** |
| Asp | **0.67** | **<0.001** |
| Ala | 0.38 | 0.12 |
| Val | **0.40** | **<0.01** |
| Pro | **0.45** | **<0.01** |
| Ile | **0.50** | **<0.001** |
| Leu | **0.42** | **<0.01** |
| Phe | 0.03 | 0.80 |
| Lys | 0.21 | 0.18 |
| Gly | -0.27 | 0.04 |
| Ser | -0.08 | 0.54 |
| Met | 0.37 | **0.01** |
| His | 0.05 | 0.74 |
| Tyr | 0.15 | 0.25 |
| Thr | -0.10 | -0.53 |

**Table S6**. Pearson correlations between TEF_cod-herring_ in each amino acid (AA) and the fat% (arcsine transformed) and Fulton’s K in cod during the deterioration period 1994-2018 (no data for herring in 1993 and 2019) and the earlier time period (1981-1994). Values in bold are lower than the Bonferroni corrected (n=15 AAs) p-value of 0.013.

|  |  | | Fat% | | Fulton’s K | |
| --- | --- | --- | --- | --- | --- | --- |
| Period | | AA | Pearson’ r | p-value | Pearson’ r | p-value |
| Deterioration period  1994-2018 | | Glu | **-0.67** | **0.002** | **-0.83** | **<0.001** |
|  |  | Asp | **-0.66** | **0.002** | **-0.78** | **<0.001** |
|  |  | Ala | -0.41 | 0.078 | **-0.61** | **0.005** |
|  |  | Val | **-0.61** | **0.005** | **-0.61** | **0.006** |
|  |  | Pro | **-0.56** | **0.012** | **-0.57** | **0.011** |
|  |  | Ile | -0.46 | 0.044 | **-0.61** | **0.006** |
|  |  | Leu | **-0.67** | **0.002** | **-0.77** | **<0.001** |
|  |  | Phe | -0.46 | 0.049 | -0.44 | 0.058 |
|  |  | Lys | -0.34 | 0.148 | -0.39 | 0.100 |
|  |  | Gly | -0.49 | 0.035 | -0.53 | 0.018 |
|  |  | Ser | -0.32 | 0.187 | -0.48 | 0.038 |
|  |  | Met | -0.28 | 0.248 | -0.30 | 0.215 |
|  |  | His | **-0.63** | **0.004** | -0.34 | 0.153 |
|  |  | Tyr | -0.29 | 0.224 | -0.42 | 0.071 |
|  |  | Thr | 0.42 | 0.076 | 0.25 | 0.299 |
| 1981-1994 | | Glu | 0.51 | 0.088 | 0.47 | 0.122 |
|  |  | Asp | 0.55 | 0.063 | 0.60 | 0.038 |
|  |  | Ala | 0.43 | 0.164 | 0.43 | 0.166 |
|  |  | Val | 0.20 | 0.542 | 0.27 | 0.400 |
|  |  | Pro | 0.27 | 0.398 | 0.78 | **0.003** |
|  |  | Ile | 0.57 | 0.052 | 0.41 | 0.180 |
|  |  | Leu | 0.54 | 0.071 | 0.48 | 0.118 |
|  |  | Phe | -0.17 | 0.604 | 0.06 | 0.850 |
|  |  | Lys | -0.06 | 0.305 | -0.06 | 0.848 |
|  |  | Gly | -0.36 | 0.248 | -0.05 | 0.885 |
|  |  | Ser | -0.30 | 0.349 | 0.14 | 0.657 |
|  |  | Met | -0.27 | 0.403 | 0.03 | 0.932 |
|  |  | His | -0.39 | 0.216 | -0.12 | 0.718 |
|  |  | Tyr | -0.58 | 0.051 | -0.19 | 0.565 |
|  |  | Thr | -0.63 | 0.029 | -0.34 | 0.284 |

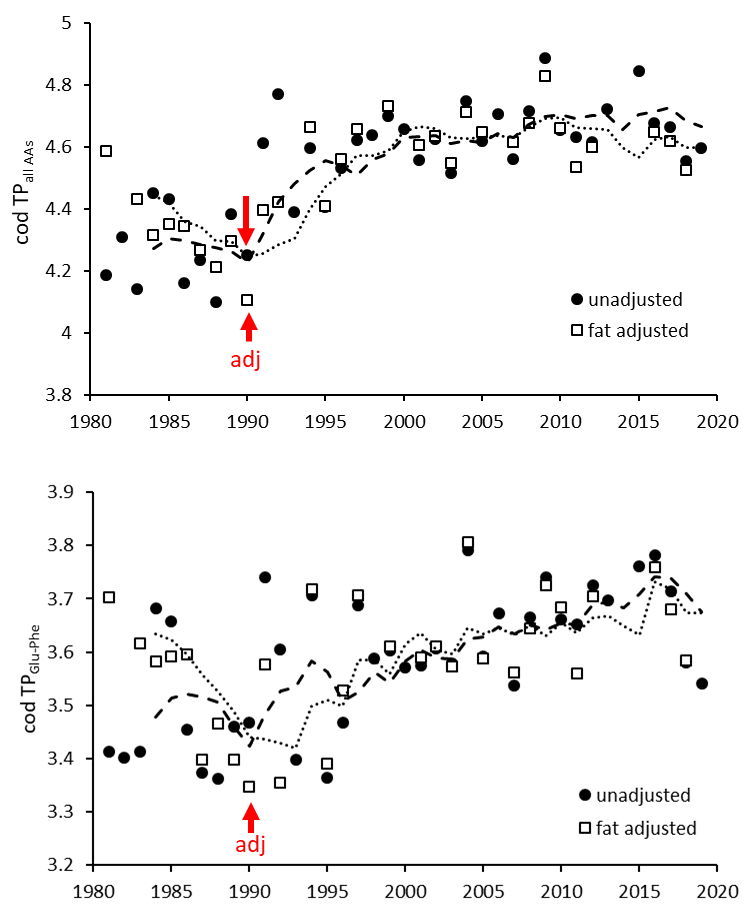

**Fig S1.** Trophic position (TP) in cod calculated using all AAs (upper) and two AAs (Glu and Phe) with and without adjustment for physiological status (fat in both cod and its prey herring, see Table 1). Compare Fig 4 where a second adjustment for hypoxia is shown. Red arrows indicate statistical breakpoints (1990, but no breakpoint was detected for unadjusted TP_glu-phe_ (Table 1 for details on time trends as assessed from Kendall-tau correlations after breakpoints).

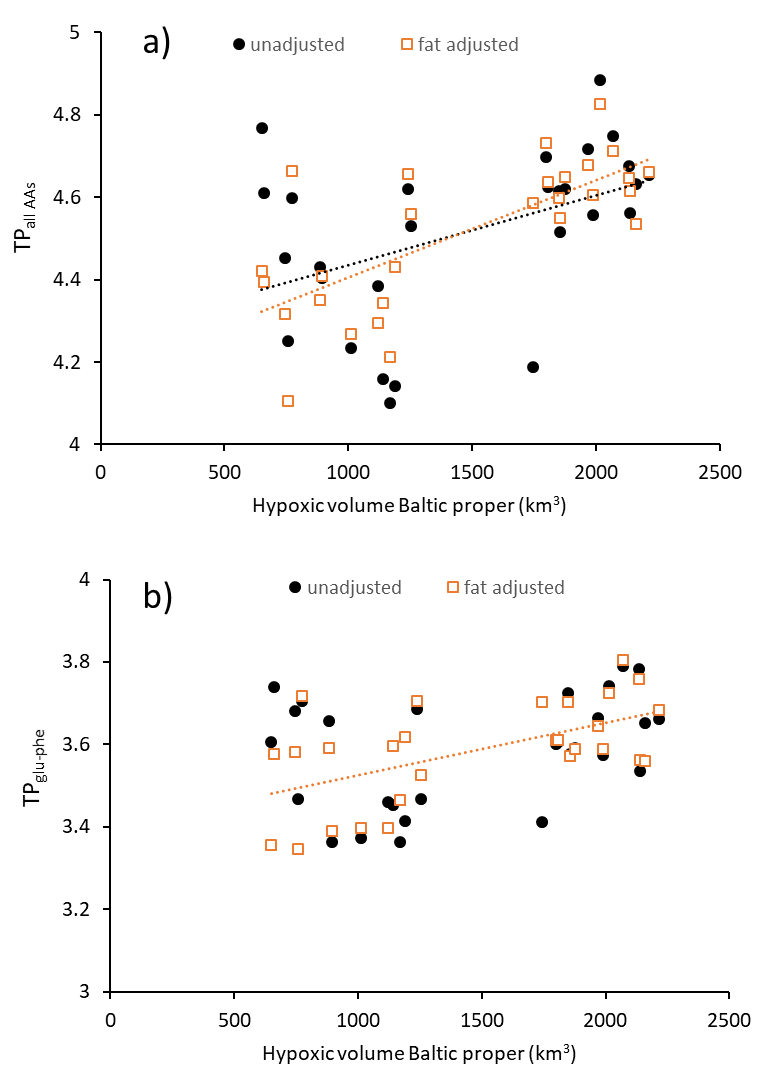

**Fig S2**. Pearson correlations (significant correlations are indicated with trend line) between trophic position (TP) calculated using all AAs (a) and two AAs (b) with hypoxic volume in the Baltic proper (data from Supplementary Table 3 in Savchuk 2018, Front Mar Sci doi.org/10.3389/fmars.2018.00095). In (a) the r-values (pearson coefficient) was r=0.44 p<0.02 for unadjusted and for adjusted data r=0.72 p<0.01. Corresponding values in (b) were r=0.33, p<0.10 and r=0.56, p<0.01.

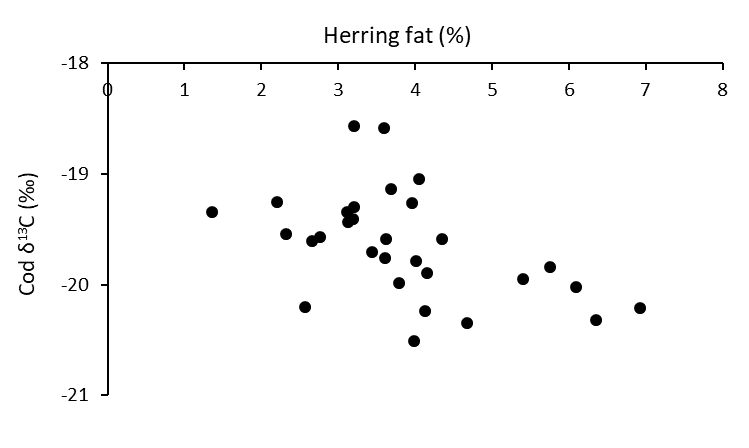

**Fig S3.** Spearman correlation (rho = -0.55, p<0.01) between herring fat (%) and cod δ^13^C during the entire time series (1981-2018).
